# Motion prediction enables simulated MR-imaging of freely moving model organisms

**DOI:** 10.1101/598524

**Authors:** Markus Reischl, Mazin Jouda, Neil MacKinnon, Erwin Fuhrer, Natalia Bakhtina, Andreas Bartschat, Ralf Mikut, Jan G. Korvink

## Abstract

Magnetic resonance tomography typically applies the Fourier transform to *k*-space signals repeatedly acquired from a frequency encoded spatial region of interest, therefore requiring a stationary object during scanning. Any movement of the object results in phase errors in the recorded signal, leading to deformed images, phantoms, and artifacts, since the encoded information does not originate from the intended region of the object. However, if the type and magnitude of movement is known instantaneously, the scanner or the reconstruction algorithm could be adjusted to compensate for the movement, directly allowing high quality imaging with non-stationary objects. This would be an enormous boon to studies that tie cell metabolomics to spontaneous organism behaviour, eliminating the stress otherwise necessitated by restraining measures such as anesthesia or clamping.

In the present theoretical study, we use a phantom of the animal model *C. elegans* to examine the feasibility to automatically predict its movement and position, and to evaluate the impact of movement prediction, within a sufficiently long time horizon, on image reconstruction. For this purpose, we use automated image processing to annotate body parts in freely moving *C. elegans*, and predict their path of movement. We further introduce an MRI simulation platform based on brightfield-videos of the moving worm, combined with a stack of high resolution transmission electron microscope (TEM) slice images as virtual high resolution phantoms. A phantom provides an indication of the spatial distribution of signal-generating nuclei on a particular imaging slice. We show that adjustment of the scanning to the predicted movements strongly reduces distortions in the resulting image, opening the door for implementation in a high-resolution NMR scanner.

## 1 Introduction

A major challenge in biological science is to relate molecular regulation at the cellular level to response and behaviour at the organism level. Knowing this relationship lies at the foundation of every disease, and indeed also in understanding the healthy organism. An experiment establishing this relationship, as schematically shown in Fig. 1, requires i) *in vivo* cellular-level detection of regulation-relevant molecules, such as metabolites and their production rates, ii) precise access to and application of biological perturbation mechanisms, and iii) an unburdened (naturally responding) organism. Clearly, even taken individually, these items are very hard to achieve in general, and correspond to major research areas in their own right.

**Fig 1.**
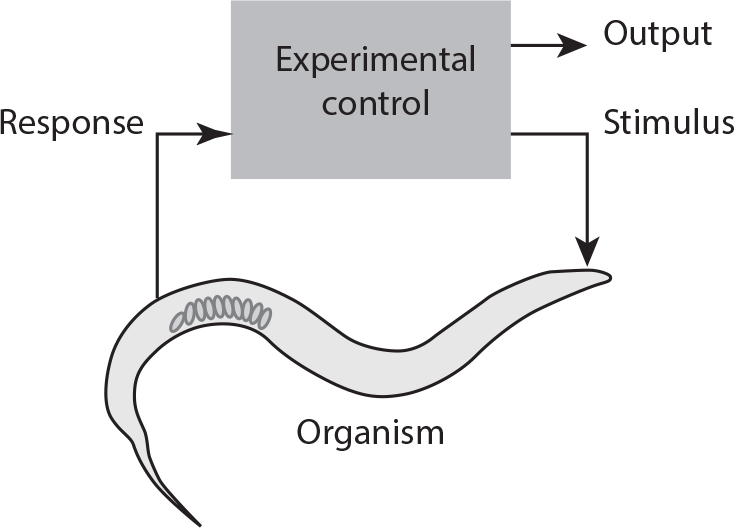
A freely moving organism is subject to a stimulus (mechanical, chemical, light, etc.) causing a response at all levels of detail (metabolomic, behaviour, etc.). The experiment designer relates the correlated output to a biological hypothesis and may adapt the stimulus.

Small organisms such as *C. elegans* are biological model organisms useful for studying many human disorders, including neurodegenerative diseases [1, 2]. Model organisms have been the mainstay of biological sciences for decades, and thus a broad knowledge base already exists starting from the genome level, through developmental cycles, and up to behavioural response to applied stress. These platforms offer the opportunity to address the connection between molecular phenotype, which can be conveniently implemented due to the very rapid yet standardized life cycle of *C. elegans*, to behaviour. What remains is the technological challenge of satisfying the three requirements for robustly linking phenotype to behaviour.

Nuclear magnetic resonance (NMR), which is a noninvasive and non-destructive technique, together with its imaging modality (MRI), is a strong candidate as the analytical method of choice, towards the ultimate goal of *in vivo* measurement of the molecular response. NMR is based on exciting the spin-active nuclei of the magnetized organism with radio frequency (RF) signals, and detecting their response via induced RF signals. Many nuclei are NMR sensitive, but *in vivo* molecular concentrations of metabolites are typically below mmol levels, requiring high sensitivity to detect. NMR is a non-ionizing technique, making it superior to computed tomography (CT) based on X-rays.

Although MRI microscopy currently only achieves spatial resolutions down to 4 µm, new techniques are partially overcoming these limits, such as the use of nitrogen vacancy centers in diamond, or the use of hyperpolarisation techniques. The encoding of the MRI signal occurs due to magnetic field gradients ∇_*x*_B over the region of interest through the Larmor relation *ω* = *γB* for the signal frequency *ω*; any gradient in *B* results in a gradient in *ω*. The gyromagnetic ratio *γ* is specific to the nucleus of interest, and is 42.6 MHz T^*−*1^ for protons. In conventional MRI, it is assumed that the organism (or object) being imaged is fixed in space, so that spatio-temporally varying magnetic fields are only due to the technical system. Any movement of the organism results in measurements from shifted spatio-temporal positions resulting in image artifacts and ensuing difficulty in data interpretation (see Fig. 2). Motion-induced artifacts are *the* challenge since spatially localized spectroscopy (required for molecular profiling) necessarily requires repeated measurements to bring the signal level above the noise.

**Fig 2.**
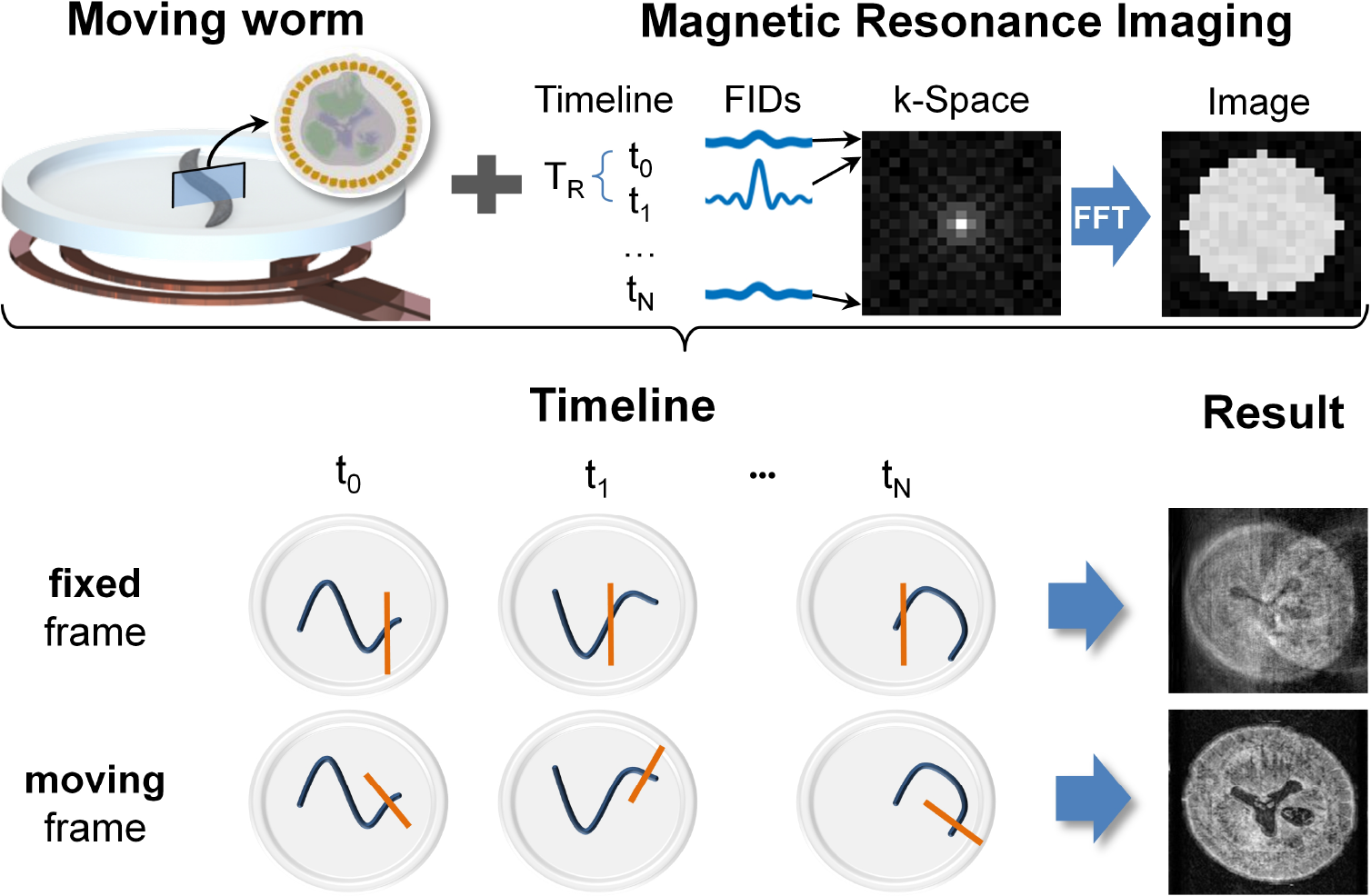
Concept of moving frame imaging: An object of interest is placed close to an NMR sensor (coil). For conventional acquisition of an MR image, *N* repetitions are required to fill the k-space image of 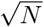 resolution, resulting in a total acquisition time of *T*_*R*_ ⋅ *N*. The time scales of worm motion and repetition time *T*_*R*_ are of the same magnitude, and with a standard Eulerian or fixed frame MR procedure, strong motion blurring occurs. To enable imaging free of blurring of a freely moving worm, the concept of a Lagrangian moving frame is required.

In clinical applications of MRI, free body motion remains a challenge and there is a concerted effort underway to i) collect the MRI data faster than the motion; ii) collect the MRI data at moments when motion is minimal (triggering); iii) track patient motion and correct the MRI data during post processing; iv) track the motion and guide the spatial encoding to reflect the instantaneous geometrical configuration [3–6]. Whilst these methods have been highly successful to control artefacts due to breathing, heartbeat, and low amplitude head movement, their assumptions for the kinematics of the underlying movement is limiting. For example, the head is motion-captured in a model that assumes rigid six-degree-of-freedom body movement involving translations and rotations (*x, y, z, θ*_*x*_, *θ*_*y*_, *θ*_*z*_) along the three orthogonal Cartesian axes. The direct translation of these techniques to MR measurements of small samples is not straightforward primarily because of the reduced size, more complex organism motion including writhing and wiggling, and rapid displacement across the detector’s sensitive region. Methods used to immobilise an organism can be considered to avoid these motion artifacts, such as clamping or freezing; however, such drastic measures typically introduce an undesired stress response into the molecular profile. The challenge of molecular measurement of non-stressed, small model organisms therefore still remains open. Given the advances in image processing, we believe there is an opportunity to address this challenge computationally.

Advances in computer vision and machine learning have revolutionized the speed and accuracy with which image analyses can be performed, and aim to reduce the need for expert knowledge e.g. in medical image analysis [7]. These technologies are entering public awareness through the automation of highly complex processes, such as trajectory generation for self-driving vehicles (road, aerial) and surveillance for public safety. This is accomplished by real-time processing of dynamic images, in which accuracy and speed are of paramount importance.

Advances in image and data processing algorithms are expected to make real-time dynamic *spectrum* imaging (achieving a hyperspectral imaging cube or hypercube) possible at all electromagnetic wavelengths. For example, recording image and spectral data over 500 × 500 pixels, with a spectral resolution of 5 nm over the visible spectrum, at 5 f s^−1^ has already been demonstrated [8]. Enormous data storage and processing power is required to perform such operations with sufficient spectral, spatial, and temporal resolution, and has motivated an increased effort in sparse imaging modalities. In the case of MRI, the situation is even more acute since data is acquired in 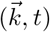-space (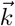 being the spatial frequency), yet the observer requires the transformed data in 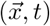 space. Hence, a large number of acquired data is associated with each final image pixel or voxel, and motion or geometrical warping during imaging will introduce errors into the reconstruction algorithms. To obtain sufficient acquired signal power per voxel volume, object tracking must be implemented in order to ensure correct spatial and spectral co-localization over time. This is especially important in MRI microscopy, which depends on accumulated sampling for sufficient image resolution. The prediction horizon will therefore be dependent on the signal acquisition time at a given frame rate and voxel spatial resolution.

In this contribution we consider the preconditions for performing MRI experiments on unburdened small organisms. We will focus our attention on the nematode *C. elegans*, mainly because it fits in best with our own efforts towards *in vivo* metabolomic profiling, but we will address the underlying problem in more generality so that it is relevant also for other organisms.

In this contribution we explore the possibility to completely remove the requirement of organism immobilisation, by providing the host observing technical system (e.g., a spectrometer or microscope) with a real time co-evolving Lagrangian coordinate system centered in the organism that provides the organism’s current center-of-gravity position and shape, paired with a robust prediction of the organism’s future position and deformed shape, in contrast to so-called motion-trackers, which mainly quantify rigid body motions and ignore strain fields.

To gain insight into how movement prediction can enhance MRI signal detection, the paper introduces:

- Data of a virtual phantom, combining high resolution electron microscope slice images [9] with conventional video-recordings of *C. elegans* moving in a Petri dish (Section 2.1);
- A new concept for location prediction in *C. elegans*, and detailed movement prediction, exploiting characteristic worm movements (Section 2.2);
- A computational platform, being able to evaluate the outcome of MR imaging with and without adapting the imaging parameters based on the movement prediction (Section 2.3).
- A measure for simulation success that compares prediction to the ‘true’ image through a similarity measure *s*_*xy*_, a technique first introduced in [10].

Using this simulation model, we demonstrate the capabilities of the motion-prediction algorithm in MRI.

## 2 Results

### 2.1 Phantom Generation

The study is based on eight 10 s duration AVI-videos (sampling frequency 12 Hz) of *C. elegans*. The recording was done in a controlled environment with a fixed camera and constant illumination over time. The worm was enclosed in a technical setup including a microfluidic channel and a Petri dish of the host system, suitable for optical recording and other real-time measurements. Typically, if natural state studies are relevant, the organism can also be provided with an optically transparent nutritional substrate such as a gel containing *E. coli* bacteria.

To simulate MR imaging of the moving worm, we artificially linked transmission electron microscope (TEM) images to the video of the worm, such that virtually scanning each location within the worm would deliver a simulated high-resolution MR-image. At each time-frame, the 1.2 mm worm was segmented into 50 slices perpendicular to its center line. This resulted in a reasonable slice thickness of roughly 24 µm. Subsequently, we assigned voxel MR signals adapted from the TEM images to each slice as follows: first, we removed the background from the TEM images and scaled them such that the dimensions of the body part in each slice were realistic. Second, we inverted the color map of the images, since the white regions of the TEM images corresponded to low density material, and therefore would appear dark in MR imaging due to the low proton signal intensity. Finally, we reduced the number of pixels of the TEM images to 64 × 64, which corresponded to an MRI in-plane resolution of approximately 1.6 µm, and then assigned the voxel signals to the virtual MR slices. Although such a high volumetric resolution lies beyond the capabilities of currently available MRI scanners, it was chosen on purpose to allow more accurate assessment of the prediction algorithm.

### 2.2 Motion prediction

#### 2.2.1 New concept

In order to adjust the gradient system of the MRI, the worm needs to be detected and its future positions need to be predicted. Therefore, we introduced a new concept of real-time image processing of the worm, suitable for any video format, regardless of color model, and also independent of background structures.

Fig. 3a shows a block diagram of the basic steps of the concept. Starting with a raw image matrix *X*_raw_, a robust estimation of the worm and its position *X*_worm_ within the video were determined. Based on the segmented worm, characteristic points such as center of gravity (COG) and position of the head region were determined. To introduce a coordinate system along the center line of the worm, a skeletonization was applied, see Fig. 3b. Assuming that body regions excluding the head would move along the center line, the worm velocity was used to predict positions *x_c,p_* of arbitrary selected points *x*_*c*_ within the worm.

**Fig 3.**
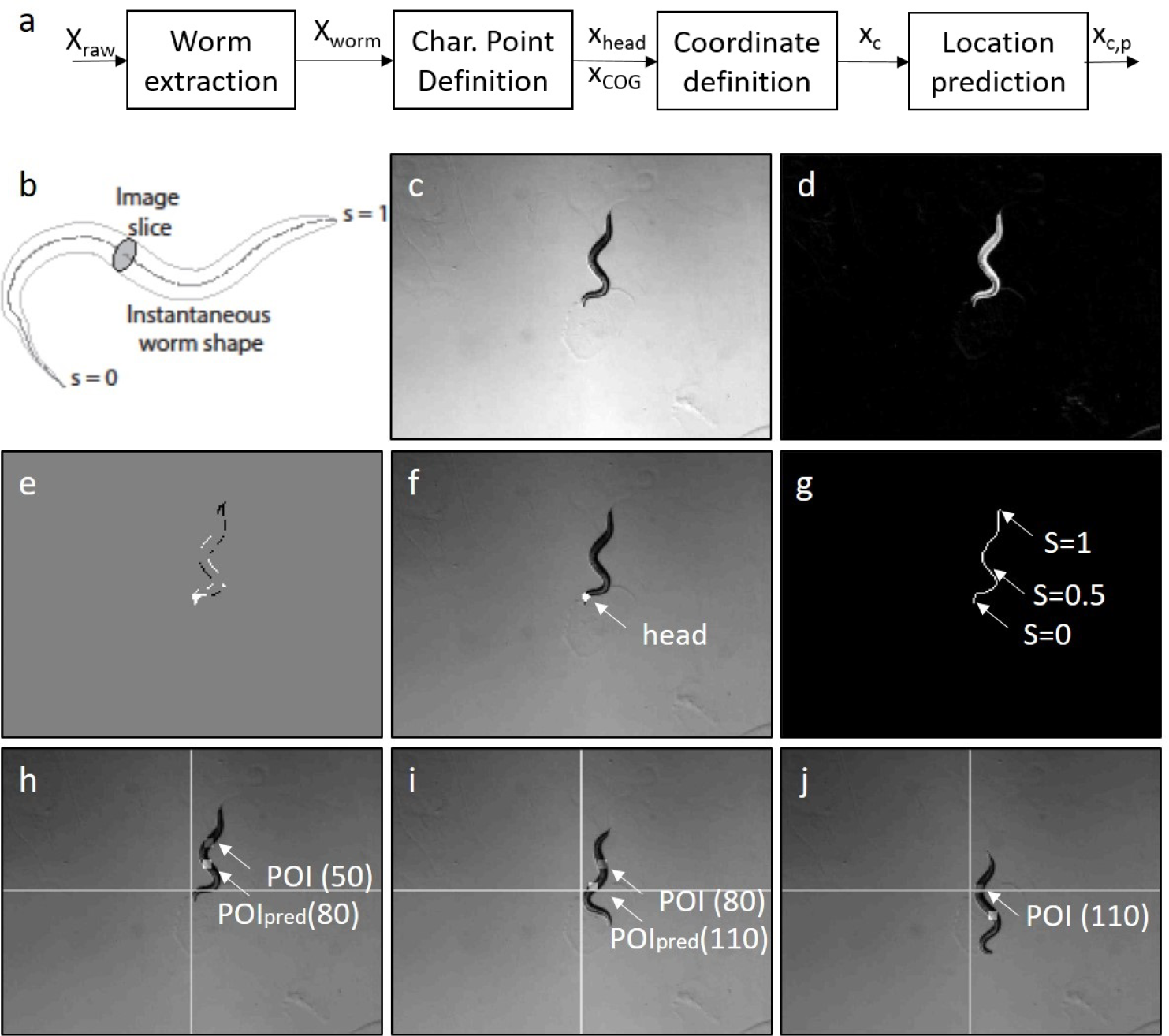
a: Concept for the prediction of defined locations within a worm, b: Parametrization of the worm, c-j: Snapshot of a sample video. c: grayscale image/video **X**^raw^[*k*], d: Image with subtracted background, e: difference image, f: overlay with worm, g: skeletonization, h-j: Point of interest (POI, gray marker) and predictions for ∆*k* = 30 (=2.5s) time samples (white marker). h: Time sample *k* = 50, i: Time sample *k* = 80, j: Time sample *k* = 110.

#### 2.2.2 Worm detection

This section briefly covers the worm detection and movement prediction. Further details are provided in the Appendix, Section 3.

The preprocessing for the worm detection aims to reduce the computational complexity and consists of a grayscale conversion (standard Matlab conversion) of the videos as well as a decrease of the resolution by a factor of 9 (Fig. 3c). For the segmentation of the worm, the background of the video is estimated and removed from the images (Fig. 3d).

The direction of motion is determined by relying on the position of the head and the COG. The COG is calculated based on the foreground pixels of the segmentation and is smoothed to obtain variations for future predictions (1st order low-pass filtering over time). The position of the head is computed utilizing difference-images (Fig. 3e) of the last 10 frames in the video. Fig. 3f shows the detected head.

#### 2.2.3 Coordinate prediction

The position of the COG moves linearly and is predicted by linear extrapolation based on its past five positions. To predict the position of an arbitrary point of interest (POI) within the worm (which can easily be identified e.g. by a mouse click in the an NMR-simulation), we use the skeleton line as the center line of the segmentation and introduce a normalized coordinate *s* along it (head: *s* = 0, tail: *s* = 1, see Fig. 3b, 3g).

Assuming that each worm segment moves according to the current shape of the worm, following its predecessor segment with the velocity of the worm^1^, the velocity and the shape is used to predict the location *s*_*c*_ of the POI after ∆*k* time steps using *s*_*c*_ = *s − v*∆*k*, (Fig. 3h-j)^2^.

The prediction quality is evaluated using an Euclidean distance. Given the prediction horizon ∆*k*, the predicted position based at time point *k* + ∆*k* is calculated (*x*_1,*c,p*_[*k* + ∆*k*], *x*_2,*c,p*_[*k* + ∆*k*]) and the distance to the true position (*x*_1,*c*_[*k* + ∆*k*], *x*_2,*c*_[*k* + ∆*k*]) is measured. The average for all time samples is termed the mean prediction error. The dependency of the prediction accuracy to the prediction horizon, as well as to the predicted position of the worm, is given in Fig. 4: The prediction error increases if the prediction horizon becomes larger. Regions near the head cannot be reliably predicted, since the head moves at a much higher frequency and in random patterns while scanning the surroundings and deciding on the moving direction. Furthermore, if the prediction horizon is set too high the predicted point lies outside the current shape of the worm (e.g for ∆*k* = 12 for all *s* < 0.3) and the topmost point (s = 0) is chosen as prediction, resulting in deviations)

**Fig 4.**
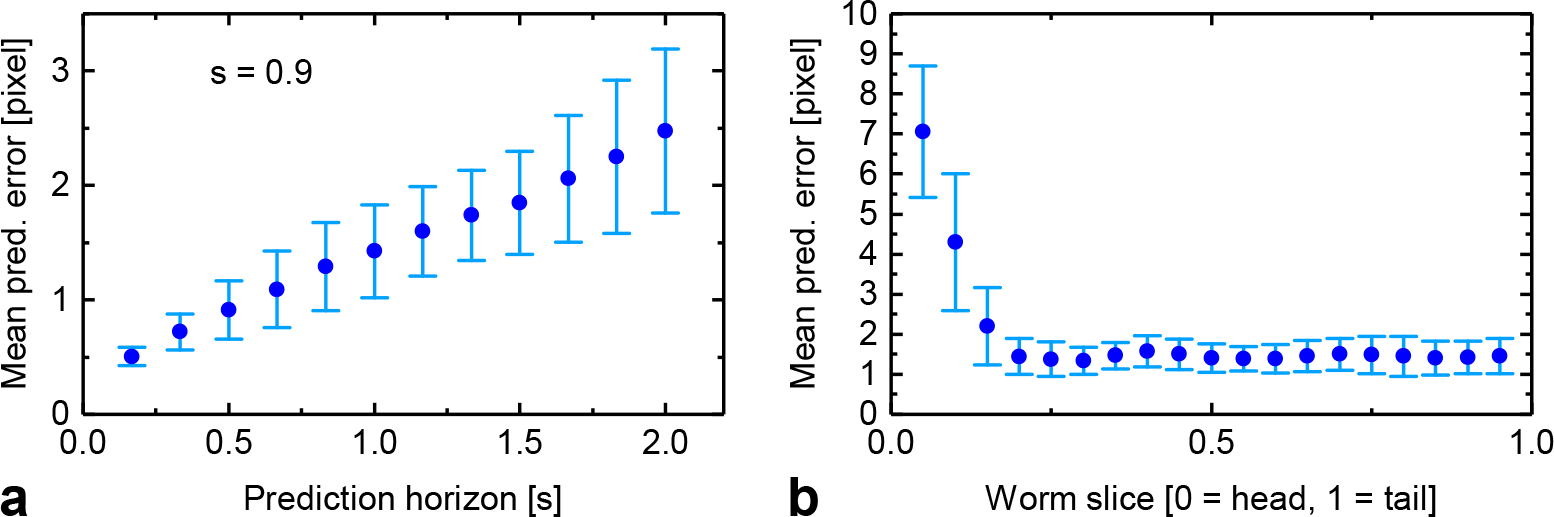
Evaluation (mean values and standard deviation) of the dataset given in Sec. 2.1. a: fixed location in the worm (*s* = 0.9), varying prediction horizons, b: fixed prediction horizon (∆*k* = 12, 1 second, varying locations, prediction horizon too high for *s* < 0.3).

For all other regions, the uncertainty of the prediction stays roughly constant.

### 2.3 MRI Simulation

#### 2.3.1 Simulation-Platform

The magnetic resonance imaging process is simulated using Matlab. The worm is assumed to reside within a strong and constant magnetic field *B* = (0, 0, *B*_*z*_) of the MRI scanner. The software mimics a simplified yet acceptably accurate MR image acquisition process via a standard gradient-echo sequence [11] that takes place during worm motion as follows:

- An image slice at a parametrised position *s* along the worm axis is defined as the image plane perpendicular to the worm axis. This slice is excited by assigning a signal intensity value for each voxel of the slice as a function of the corresponding phantom TEM image. The slice excitation is followed by a magnetic resonance phase encoding step, in which each voxel of the selected slice is given a phase proportional to its position along one cross-sectional axis *ξ* of the slice using an applied magnetic field gradient ∇_*ξ*_*B*_*z*_. After a time delay TE/2, where TE (echo time) is the time from excitation to the center of the MR signal (echo), a frequency encoding gradient is applied, whereby each voxel of the slice is assigned a frequency proportional to its position along an axis *η* perpendicular to the phase encoding axis (so that *ξ · η* = 0). The superposition of all voxel RF signals are simultaneously detected by a virtual coil. This so-called echo is recorded at instant TE, resulting in one line of *k*-space. The detection system is assumed to have a uniform spatial sensitivity.
- After a time delay TR (repetition time), which is the time it takes to repeat the sequence in order to acquire a new line of the *k*-space, the algorithm calculates the anticipated new position and orientation of the selected slice (as the worm would have already moved to a new position). Now the slice at this new position is excited, phase encoded, and frequency encoded, resulting in a new line of *k*-space being filled.
- The entire process is repeated until all the lines of *k*-space (in our case 64 lines) are complete.

Once the imaging procedure is complete, the program reconstructs the anticipated MR image from the k-space data via Fourier transform.

#### Algorithm 1 Gradient-echo MR imagings

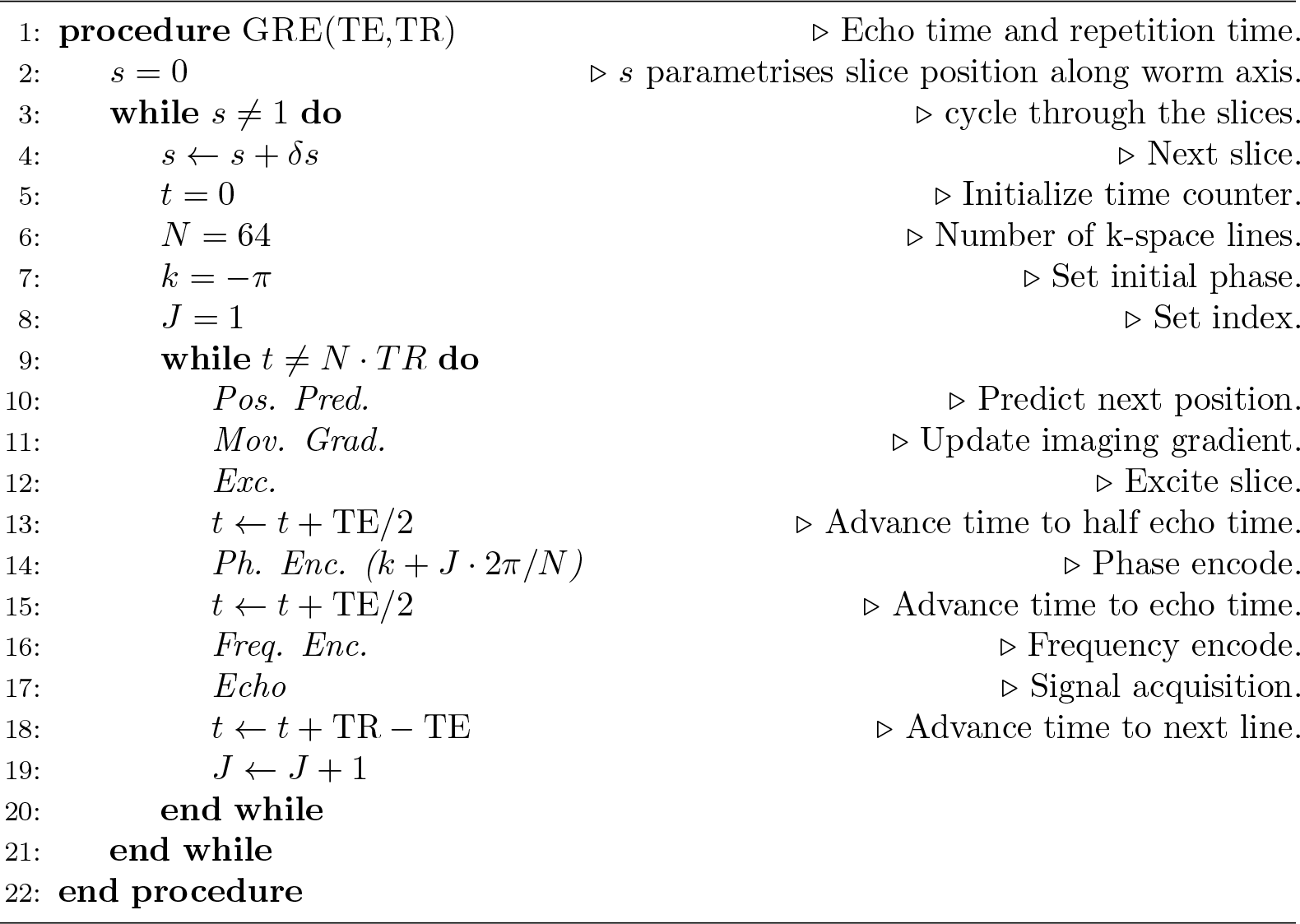

The simulation is integrated into an interactive graphical user interface (GUI) to enable the user to set and change the imaging parameters easily and execute the processing without any programming skills (Fig. 5).

**Fig 5.**
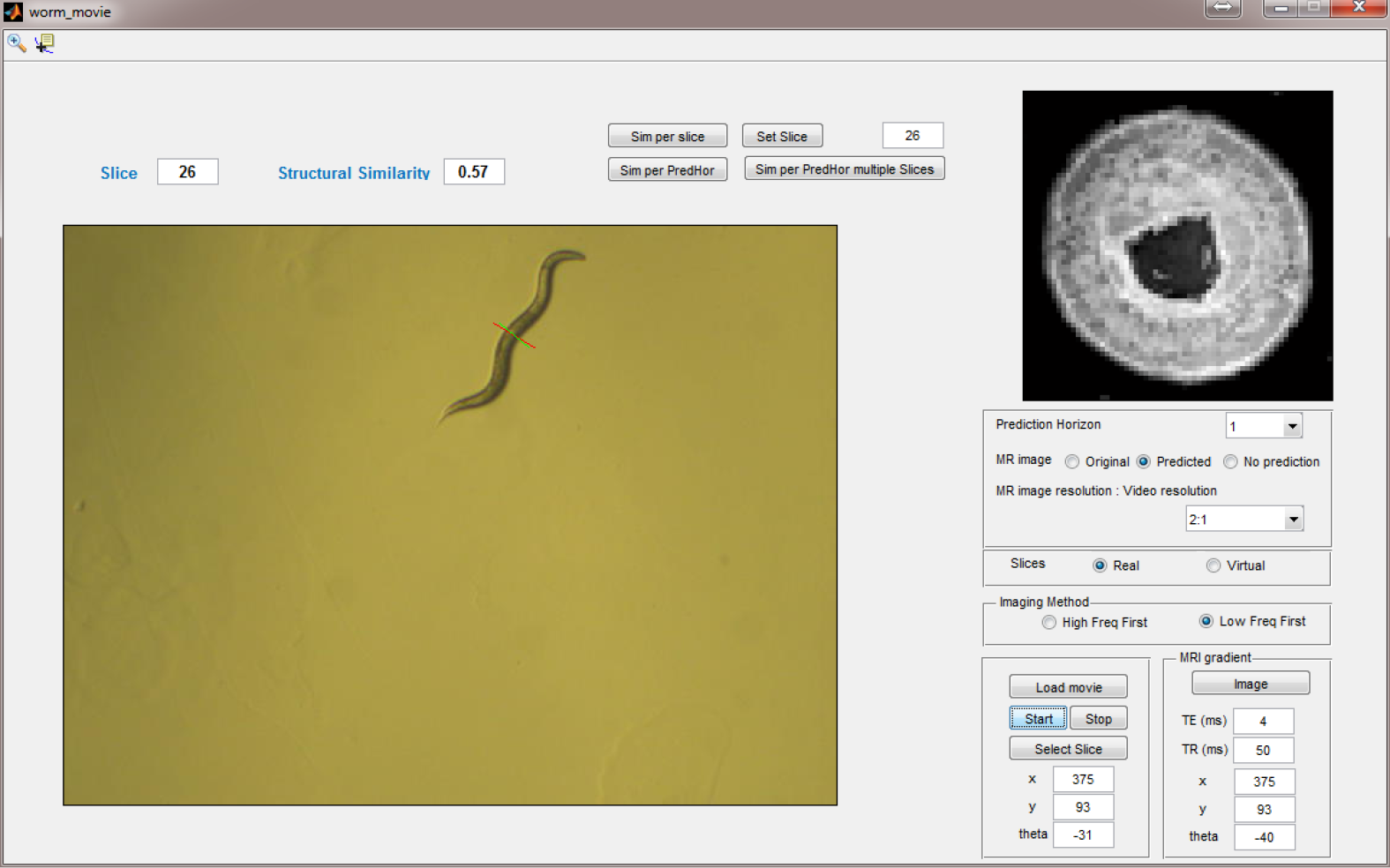
Graphical user interface of a Matlab program that simulates the MR imaging experiment of a moving worm. The microscope movie of the worm is shown to the bottom left, and emulates real-time observation. The slice position is also shown. The top left shows the phantom MR image of the worm at the position of the slice.

The simulated MRI signals can be chosen by the user and are either MR-like signals translated from real transmission electron microscopy (TEM) images, or any virtual MR-like images provided by the user.

To start the simulation, an arbitrary slice of interest (with coordinate *s*) within the worm’s body is selected. For this point, the simulated MR image of a slice without movement is shown. In the display, the slice position *s* is denoted by a red line perpendicular to the center line of the worm, while the predicted position to which the virtual imaging gradient coordinate is set is denoted by a green line.

Furthermore, the software allows the user to set the TR and TE imaging parameters, and to choose the prediction horizon (a value between 1 and 10 frames), which refers to the number of frames ahead for which the predicted worm position will be calculated. Our algorithm does not correct for the worm motion occurring during recording of a line of k-space. This is equivalent to the assumption that the MRI pulse sequence is based on very short echo times, which is possible, but taken at the expense of increased acquisition bandwidth and thus reduced signal-to-noise ratio (SNR).

#### 2.3.2 Simulation paradigms

We performed three MRI simulations using the optical video dataset from Section 2.1 to:

1. confirm that the prediction enhances the MR imaging of the moving worm in general (Simulation 1);
2. measure the efficiency of the prediction algorithm as the desired resolution of the MR images increases in comparison to the resolution of the optical video used for the prediction (Simulation 2); and
3. measure the effect of an increased prediction horizon (the number of frames ahead for which the position is predicted, Simulation 3).

In all the simulations, the repetition time (TR) was set to 83 ms, which corresponds to the frame rate of the optical video on which the prediction was based.

We evaluate the simulation by quantifying the structural similarity *s*_*xy*_ ∈ {0, 1} ∈ ℝ between the true image and simulated image as introduced in [10]. Identical images return *s*_*xy*_ = 1, whereas structural inequality delivers *s*_*xy*_ = 0. A detailed description is provided in Appendix 3

#### 2.3.3 Body position (Simulation 1)

In Simulation 1, three slices (slice definition from Sec. 2.1, first slice from the head (*s* = 0.02), second from the middle body (*s* = 0.5), third from the tail (*s* = 1) of the worm) are selected and an MR imaging simulation is performed (prediction horizon ∆*k* = 1 (83ms)).

The results are illustrated in Fig. 6a, each row shows the true slice on the left, the simulated MR image based on the prediction algorithm in the middle, and the simulated image when no position prediction is involved on the right.

**Fig 6.**
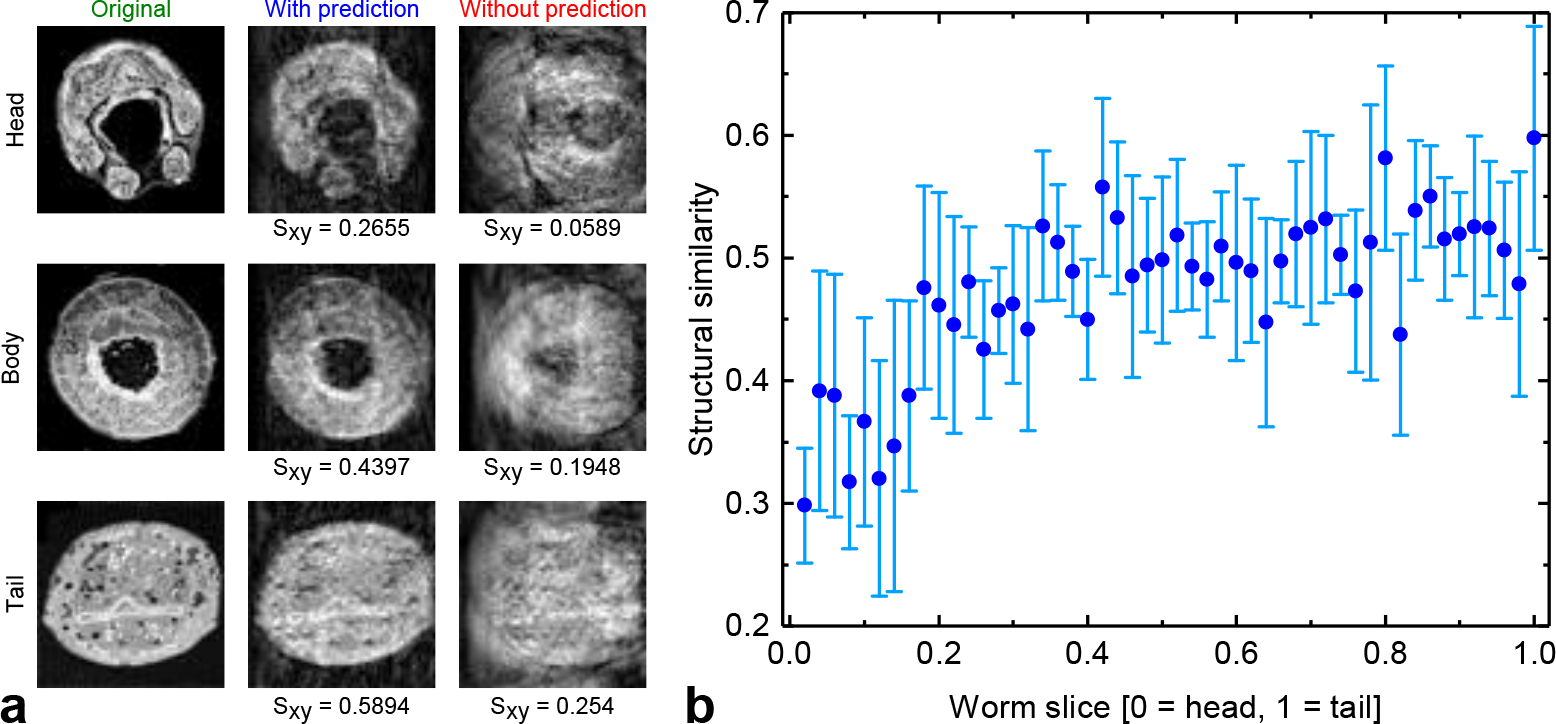
a: effect of position prediction on the quality of the MR images in the head (*s* = 0.02), body (*s* = 0.5) and tail (*s* = 1) regions. The prediction horizon in this case is 83 ms. b: dependence of the MR imaging quality (structural similarity) on the position of the slice. The figure displays the simulation results from eight videos of different worms.

Clearly, the prediction algorithm significantly reduces the motion artifacts that would otherwise occur if the gradient system did not follow the worm as it moves. Moreover, we observe that prediction quality varies along the worm’s body. More specifically, the prediction performs better for the slices from the body and tail when compared with prediction of the head. This is axiomatic, since the worm rapidly moves its head laterally whilst scavenging for nutrition, thus the motional entropy of the head is higher (see also Fig. 4). Fig. 6b shows the simulation results of eight videos of different worms. The abscissa represents the slice position starting from the head (s = 0) to the tail (s = 1), while the ordinate shows the structural similarity between the prediction-based MR images and the original slices. To very good agreement with Fig. 4, Fig. 6b shows that the quality of the prediction-based MR images are higher for slices from the body and tail than the images taken near the head where the prediction uncertainty is usually higher.

#### 2.3.4 Image resolution (Simulation 2)

In Simulation 2, a slice from the middle body section of the worm (*s* = 0.72) was selected and an MRI simulation was performed for different resolution ratios (= MR image: optical video).

Decreasing the resolution can be used to speed up image processing, and thus decreasing the prediction horizon, if needed. The ratios of 4:1 and 2:1 are heuristically chosen, for which each pixel in the optical video respectively corresponds to 4 and 2 pixels in the MR image. Fig. 7 demonstrates the results of this simulation - the rows correspond to the different resolution ratios while the columns depict the original slice, the image with prediction, and the image without prediction. The figure shows that the efficiency of the prediction declines as the desired resolution of the MR image increases (pixel size decreases), or alternatively, the optical resolution should be higher or at least equal to the desired MR resolution for a high quality prediction-based MR image. Indeed, the commercial optical imaging solutions can easily meet such demands of high resolution; however, upon implementation, any increase in number of pixels will be at the expense of the prolonged processing time needed for prediction. Nevertheless, considering the large potential for parallel computing co-processors, good compromises can be found.

**Fig 7.**
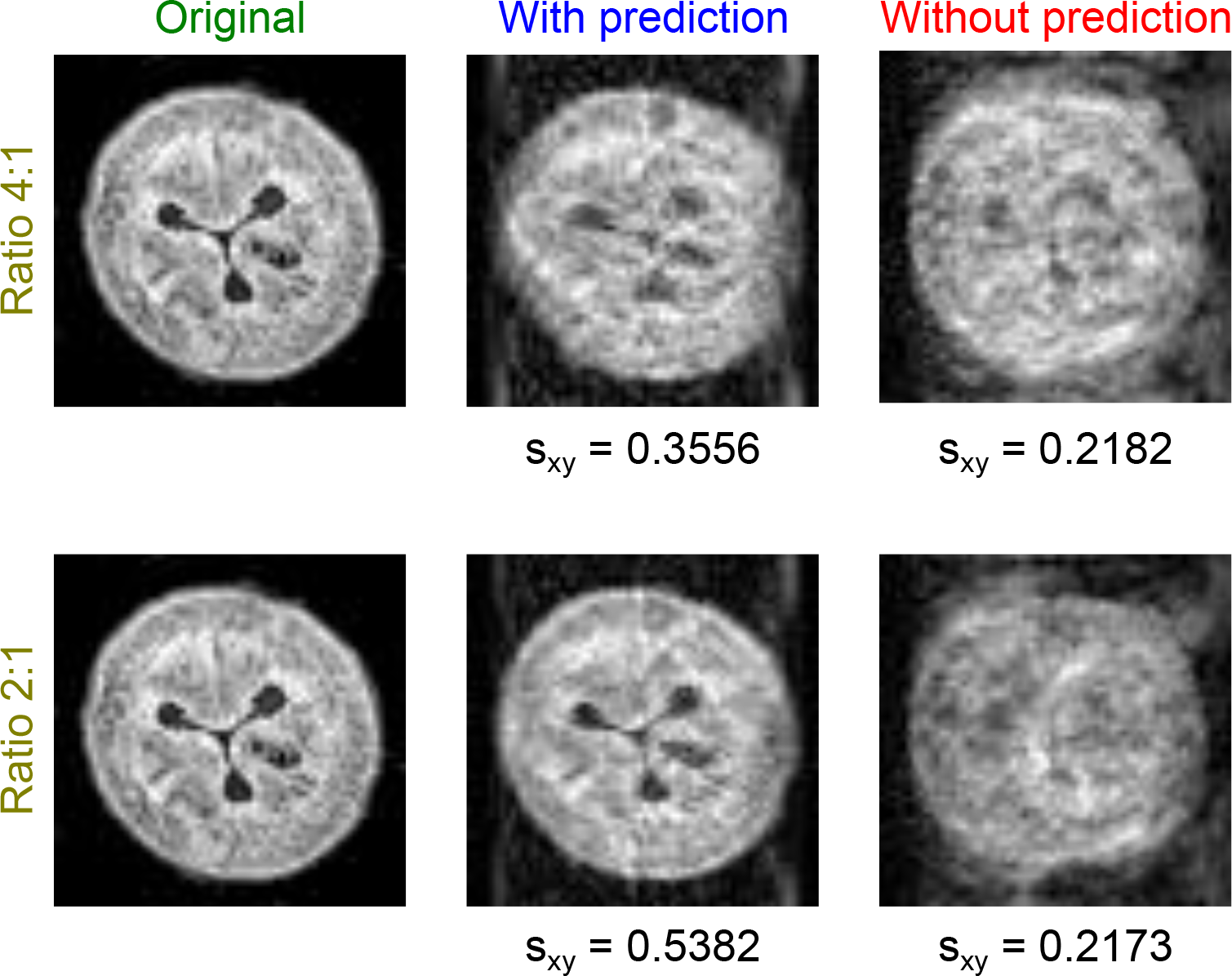
Effect of decreasing the ratio of the MR image resolution to the optical video resolution on the imaging quality for a prediction horizon of 1. The rows show the results for MR image:video resolution ratios of 4:1 and 2:1 respectively. Each row displays, from left to right, the reference slice, the MR image with position prediction, and the MR image without position prediction.

#### 2.3.5 Prediction horizon (Simulation 3)

Simulation 3 varies prediction horizons for one slice (*s* = 0.52). Fig. 8 illustrates the results of this simulation: Fig. (1a - 10a) are the prediction-based simulated images for prediction horizons of ∆*k* 1 to 10, respectively. In contrast, Fig. (1b - 10b) display the results when prediction is not in action. Because only a few lines of the k-space are collected from the correct slice (depending on TR and the speed of the worm), the images in this figure exhibit a noticeable decrease in quality as the prediction horizon increases, leading to the conclusion that one should, whenever possible, minimize the prediction horizon. Of course, in an actual hardware realization of the system, the choice of the prediction horizon will be bounded by the speed of the image acquisition and processing units.

**Fig 8.**
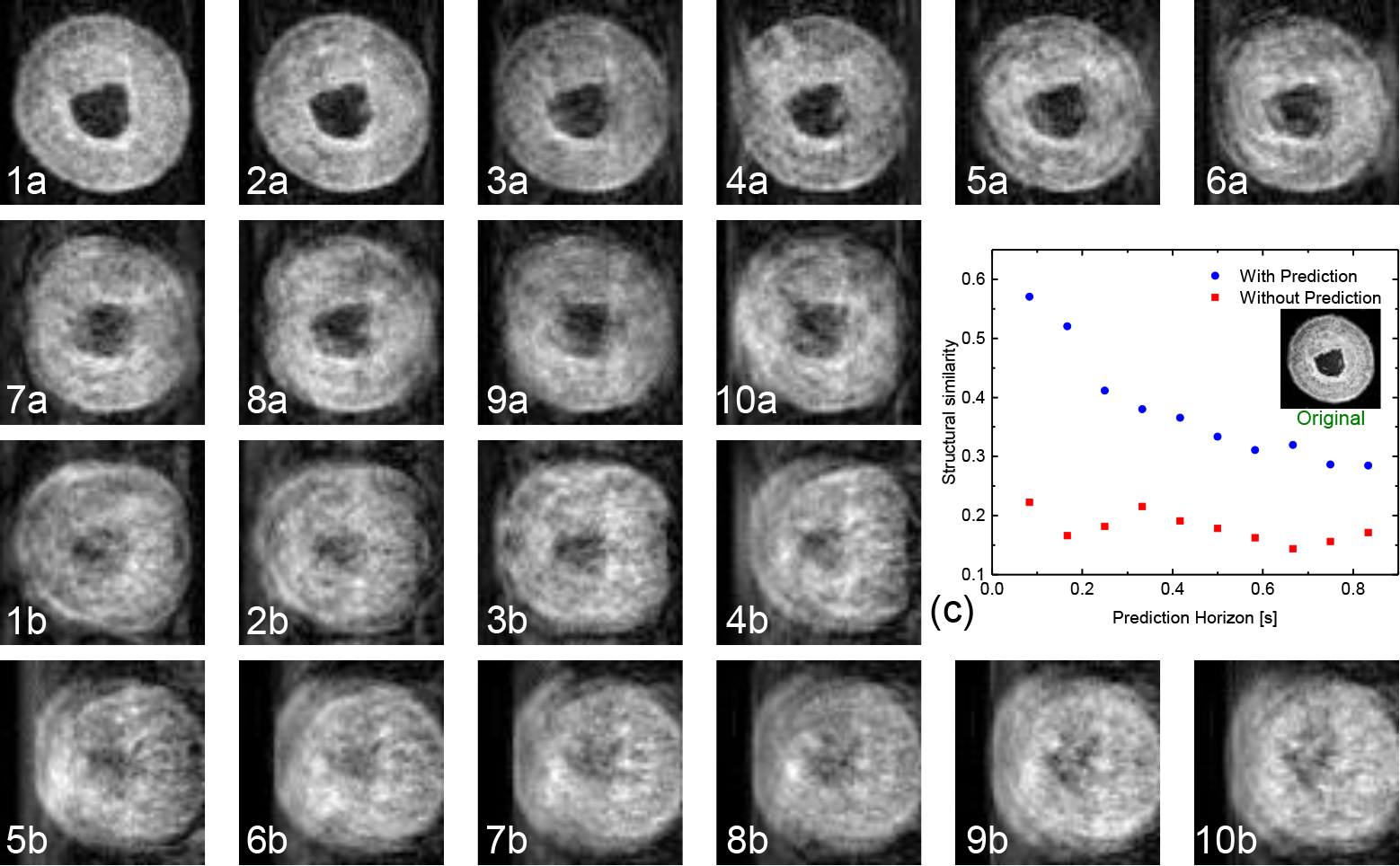
Effect of enlarging the prediction horizon on the quality of the imaging (*s* = 0.52, see Fig 6). The images (1a-10a) show the reconstructed MR images based on the prediction algorithm for prediction horizons from 1 to 10 frames. The images (1b-10b) show the reconstructed MR images when no prediction is involved and when the initial position of the imaging gradient is set to the position of the slice after 1 (83 ms) to 10 (830 ms) frames respectively. (c) The similarity measure of the simulated image versus the prediction horizon.

The effect of increasing the prediction horizon is described quantitatively in Fig. 8c by measuring the structural similarity between the original slice and the simulated image for both cases with and without prediction. The results confirm the variation in predictability along the length of the worm. The prediction horizon degradation is linear for the rear two-thirds of the worm, but falls off more rapidly for the head section, as expected from Fig 4.

Moreover, a statistical assessment of the effect of the prediction horizon on the quality of the MR imaging was done: Regarding simulation results of four slices along the worm using the given eight videos it can be shown that the accuracy of prediction and thus the MRI quality decays with the increased prediction horizon but is in mean for all parameter combinations roughly three times better than without prediction. Results are shown in the Appendix, Fig. S1.

## 3 Discussion

The results demonstrate that our prediction algorithm can markedly improve MR image quality of arbitrarily moving and deforming objects. The complete processing pipeline (including functions which will not be used in real-time processing, e.g. rotation, prediction of COG, bounding boxes etc.) takes 76 ms per frame on an average Laptop PC (core I7, fifth generation, 16 GB RAM) and has various possibilities for optimization (modern hardware, software parallelisation, hardware-filtering etc.). Decreasing the resolution of the optical video offers an additional option to reduce processing time up to a factor of 8 without significant loss of quality. Based on the current implementation, real-time processing is already possible for sample frequencies smaller than 13 Hz and application of the algorithm to a real MRI system is feasible.

For the predictive information to be useful, the characteristic time of sample motion *t*_*motion*_ and predictive accuracy needs to be determined. This can be done by recording a freely moving sample with the desired optical setup followed by an image analysis routine as described in this report. Models of the motion kinematics can be tested in order to maximize the prediction horizon given a user-specified predictive accuracy threshold. The threshold can be selected based on the expected error introduced within a single voxel with pre-selected dimensions. The predictive horizon together with the calculation time *t*_*pred*_ then can be used to determine if the sample is ‘MR imageable’, i.e. by comparing the timescales to those required for MRI. For example, in this report it was observed that a prediction horizon of 83 ms requiring a calculation time of 76 ms yielded a predictive accuracy of approximately 57 % as indicated by the structural similarity measure of the prediction-based MRI simulation, Fig. 8c. On the other hand, a prediction horizon of 830 ms requiring a calculation time of 76 ms resulted in a prediction accuracy of approximately 30 %.

With the timescales of motion and predictive calculation defined, one must evaluate whether MRI is possible by comparing to the instrumental timescale.^3^ The shortest relevant timescale is the time between *spatial* encoding steps, which in the case exemplified here is the repetition time *TR* (*TR* includes *TE* and the data acquisition time). It is during *TR* that prediction and hardware adjustments must be done prior to the subsequent spatial encoding step. In the MRI simulations described here, *TR* was 83 ms (including a *TE* of 4 ms) while *t*_*pred*_ was 76 ms. Given the organism motion and hardware/experiment timescale regimes, one can estimate the potential for sample imaging with correction, summarized as follows:

- *t*_*motion*_ < *TR*: the object is not MR imageable without motion artifacts. Conditions to slow the natural motion of the sample should be identified and implemented.
- *TR* < *t*_*motion*_, *t*_*pred*_: the object is MR imageable. Careful choice of *TE* and *TR* must be done so that the prediction calculation is complete before the next spatial encoding period. This places a restriction on the types of contrast that can be implemented.
- *t*_*pred*_ < *TR* < *t*_*motion*_: the object is MR imageable. There is no restriction on the contrast weightings that can be implemented.

To further improve the predictive quality, it is important to have kinematic models appropriate to the organism to be imaged. *C. elegans* is a convenient model for this reason given the advanced studies about behavioural phenotypes [12–14] and motion decomposition using so-called Eigenworms [15]. Extension of these models to organisms featuring similar motion characteristics should be straightforward (i.e. oscillation/undulation along the long axis - worms, snakes, swimming fish). As the kinematics becomes more complicated and/or sporadic, it will become necessary to introduce a method to guide the organism in order to introduce a predictive nature to its motion (food source, temperature gradient, etc.). Alternatively, shorter prediction horizons can be targeted together with faster calculation algorithms [4, 16–18] and MR imaging sequences, most likely at the expense of image resolution.

## Acknowledgements

We sincerely thank Dr. David H. Hall of Albert Einstein College of Medicine in New York for providing the high resolution images of *C. elegans* that we used as phantoms for high resolution NMR images. These lent our images more realism. The original images are from the MRC/LMB *C. elegans* Archive now curated by the Hall lab. These were generously donated by John White and Jonathan Hodgkin to the Hall lab. Support for the *C. elegans* Archive and for WormAtlas [9] comes from NIH OD 010943 to D.H. Hall. MJ, NM, EF, NB and JGK acknowledge financial support from the European Research Council (ERC) under grant 290586 NMCEL.

## Supporting information

### Motion prediction

This section describes the principal structure of the algorithm. Free parameters are chosen heuristically for the given dataset. Raw videos are streams of RGB images with *m* rows and *n* columns at each frame *k* = 1*, …, K*. Indices *i, j* denote *x*- and *y*-position in the image, and index *r* denotes the color:

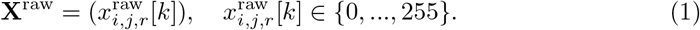

We use a standard MATLAB conversion to convert the images to grayscale and keep only 1*/z*^2^ of all pixels to speed up subsequent performance (every *z*-th line, every *z*-th column is kept, here *z* = 3, see Fig 3):

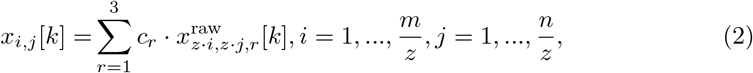

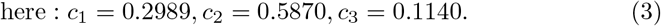

To segment the worm, a simple threshold segmentation will be erroneous due to illumination effects, noise, and dark corners. To be independent of illumination effects, we generate a background image excluding the worm by calculating at each pixel position the 75th percentile^4^ along the time scale 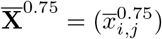. The obtained background estimation is smoothed by a closing step (structuring element **B**: DISC, *r* = 3):

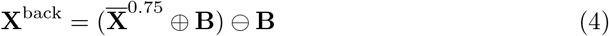

where the operators ⊕ and ⊖ respectively denote the dilation and erosion. To obtain the worm segment, we subtract the background estimation from each image frame (Fig 3d).

As a threshold for the segmentation of the worm, we choose the 1st percentile 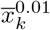 of all *m ⋅ n* pixels in the difference image at every time step (assuming that the worm covers an area of less than 1% of the image). We then discard all remaining objects except the biggest one. As an outcome, we obtain image frames **X**^worm^[*k*] of the segmented worm.

Within the worm segment, the direction of the worm needs to be identified. This can be done by estimating the position of the head and the center of gravity (COG). We calculate for each time sample *k* the COG based on the pixels in the binary image **X**^worm^[*k*]. The found values 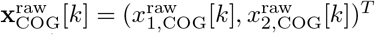 are then smoothed (1st order low-pass filtering over time) to obtain variations for future predictions. Due to quantization sometimes the COG does not move, therefore prediction algorithms might fail. The resulting time series is denoted **x**_COG_[*k*] = (*x*_1,COG_[*k*]*, x*_2,COG_[*k*])^*T*^.

To find the head, the moving parts of the worm need to be identified. Observations show that the head is always moving left/right, the tail does not. We calculate an optical flow image which contains the difference of two succeeding segmentations

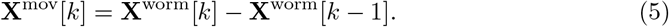

We use the integer values to distinguish between pixels where the worm moved to (value 1), and those, where it came from (value −1). For each time sample, we then sum up the last 10 optical flow images **X**^mov^[*k*] (this could also be done by simply subtracting images with ∆*k* = 10, discard all negative values (body moved away), apply an opening (=denoising and deletion of pixels along the complete body). The resulting values represent the optical flow of the head and by applying the same COG approach as before the position of the head is defined. The resulting time series is denoted 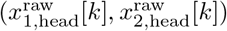.

If *k* = 1 (video just started), we set the head estimation to the COG, and if no head estimation can be calculated (e.g. due to missing movement), we retain the last one. As a final step, we smooth (low-pass filtering over time) the estimated head position values. The resulting time series is denoted (*x*_1,head_[*k*]*, x*_2,head_[*k*]).

The center of gravity is predicted by linear extrapolation (quadratic functions do not work!) according to

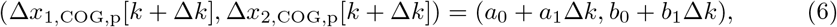

*a*_*i*_, *b*_*i*_ are estimated based on the last five values of (*x*_1,COG_[*k − j], x*_2,COG_[*k − j], j* = 0*, …*, 4) (ordinary least squares). Predictions are very sensitive, therefore we low-pass filter the found (*x*_1,COG,p_[*k* + ∆*k*], *x*_2,COG,p_[*k* + ∆*k*]) values over time.

The segmented worm is filtered with a Gaussian convolution filter (to remove noisy edges), and the segment is thinned to a line (skeletonization, see Fig. 3g).

To predict an arbitrary position within the worm (which can be identified by a mouse click later-on), we introduce a normalized coordinate system along the skeleton line of the worm (0: head, 1: tail). A click point (*x*_1*,c*_[*k*], *x*_2*,c*_[*k*]) at time sample *k* can then be transformed into a percentage value *s*[*k*] ∈ [0, 1] defining the relative position of the click within the worm. For past time samples, the Cartesian coordinates of the same *s*-value are calculated and the velocity of the click point along the line coordinate system is calculated and filtered.

Assuming that the shape of the worm stays roughly the same^5^, the velocity along the shape is used to predict where the click point will be located after ∆*k* time samples. The predicted points are filtered as well. The resulting time series is denoted (*x*_1*,c,p*_[*k*], *x*_2*,c,p*_[*k*]).

### Similarity measure

We compare the simulation outcome to the true image by the structural similarity measure that was introduced in [10]. This measure assesses the similarity between images *X* and *Y*, based on the luminance *l*, the contrast *c*, and the structure *st*. Using *µ* as the brightness mean, *σ*^2^ as the brightness variance, and *σ_xy_* as the brightness co-variance, similarity is calculated according to

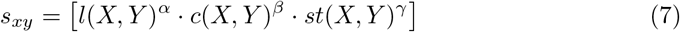

where

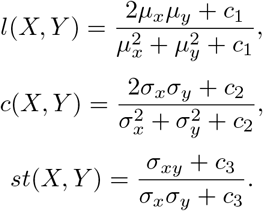

*α*, *β*, and *γ* are weights of the luminance, contrast, and structure respectively. Within this paper we assumed an equal effect of the three terms, thus *α* = *β* = *γ* = 1.

*c*_1_, *c*_2_, and *c*_3_ are small real non-negative variables used to stabilize the division when the denominator is small. The default values of these constants are *c*_1_ = (0.01 * *L*)^2^, *c*_2_ = (0.03 * *L*)^2^, and *c*_3_ = *c*_2_*/*2 with *L* being the dynamic range of the images. *S*(*X, Y*) takes values between 0 and 1, where 0 corresponds to no similarity, while 1 represents a complete similarity between the images.

### Quality of MR imaging in dependence of prediction horizon

Fig. S1 shows the statistical assessment of the effect of the prediction horizon on the quality of the MR imaging. The data in this figure is taken from the simulation results for four slices along the worm and from eight videos of different worms. The figure shows that the prediction algorithm works quite well for all worms, and the results exhibit a high degree of resemblance. Moreover, and as expected, the results demonstrate how the accuracy of prediction and thus the MRI quality decays with the increased prediction horizon.

---

**Fig S1.**
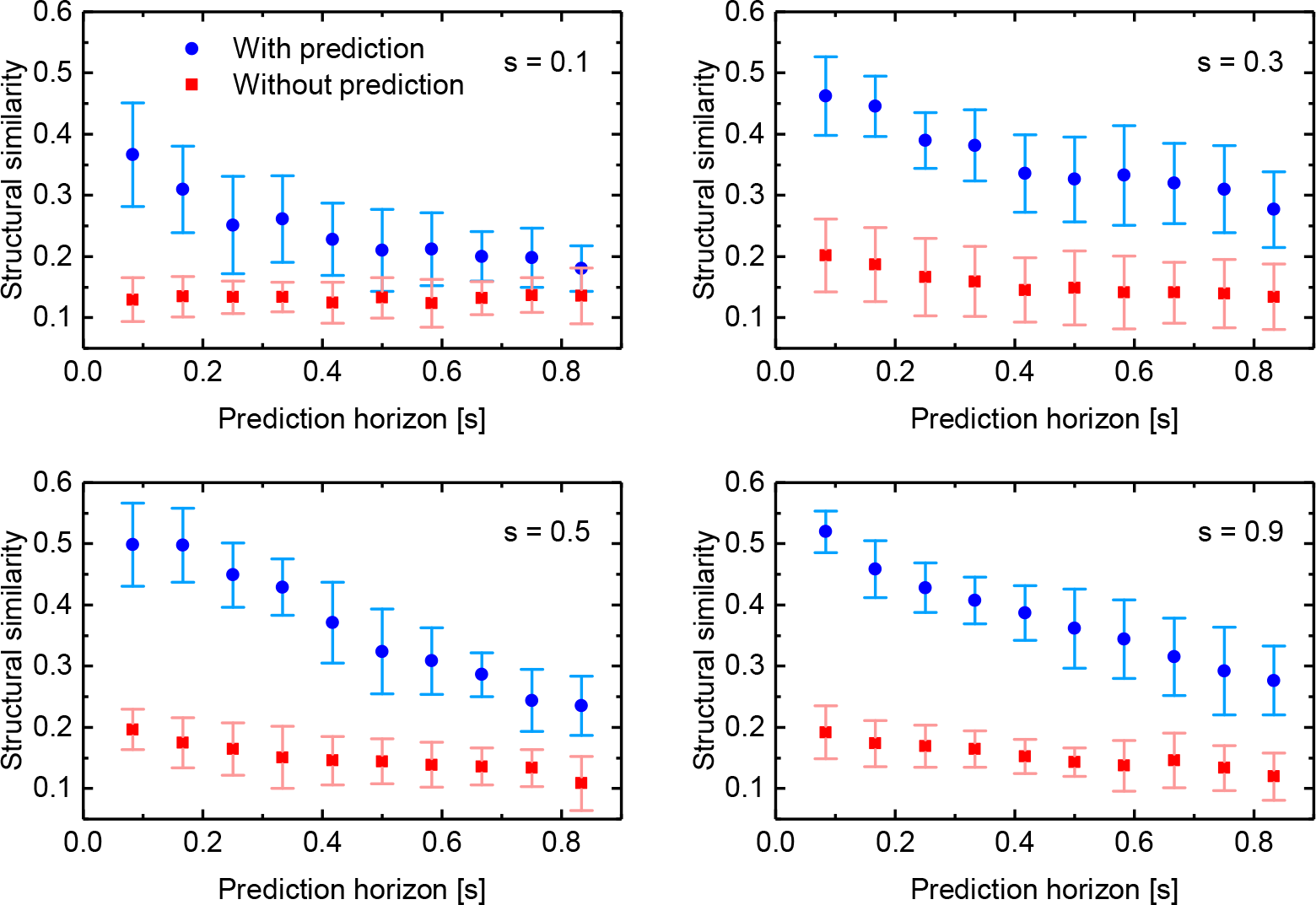
Structural similarity versus prediction horizon for slices (s = 0.1, s = 0.3, s = 0.5, and s = 0.9) for eight videos.

## Footnotes

1 The assumption is not perfectly valid, but it significantly simplifies processing and delivers reasonable results (see also Fig. 4).

2 If the prediction horizon is chosen too high, negative *s*_*c*_ are avoided by setting the prediction to the topmost point (*s* = 0).

3 Only a standard cartesian MRI sequence is considered. The authors acknowledge the existence of more advanced sampling schemes, but these are outside the scope of the present discussion.

4 The value delivers the brightness of the background pixel, if the worm does not occupy a pixel for more than 25% of the time.

5 The shape does not stay exactly the same, however this assumption simplifies processing and causes only small errors.

